# A data-driven Boolean model explains memory subsets and evolution in CD8+ T cell exhaustion

**DOI:** 10.1101/2023.03.13.532500

**Authors:** Geena V. Ildefonso, Stacey D. Finley

## Abstract

T cells play a key role in a variety of immune responses, including infection and cancer. Upon stimulation, naïve CD8+ T cells proliferate and differentiate into a variety of memory and effector cell types; however, failure to clear antigens causes prolonged stimulation of CD8+ T cells, ultimately leading to T cell exhaustion (TCE). The functional and phenotypic changes that occur during CD8+ T cell differentiation are well characterized, but the underlying gene expression state changes are not completely understood. Here, we utilize a previously published data-driven Boolean model of gene regulatory interactions shown to mediate TCE. Our network analysis and modeling reveal the final gene expression states that correspond to TCE, along with the sequence of gene expression patterns that give rise to those final states. With a model that predicts the changes in gene expression that lead to TCE, we could evaluate strategies to inhibit the exhausted state. Overall, we demonstrate that a common pathway model of CD8+ T cell gene regulatory interactions can provide insights into the transcriptional changes underlying the evolution of cell states in TCE.

## INTRODUCTION

CD8+ T cells play a crucial role in anti-tumor immunity, but they often undergo T cell exhaustion (TCE) limiting their anti-tumor effect to clear antigens on tumor cells. Exhausted CD8+ T cells are a group of dysfunctional T cells that are present in chronic infections and cancer^1,2^. An important characteristic of these exhausted T cells that persist during continuous antigen stimulation is upregulation and sustained expression of the inhibitory receptor, programed cell death protein 1 (PD1), which is also a marker of T cell activation^3,4^. Similar to other immune cells, exhausted T cells are heterogeneous and include intermediary subsets of T cell states with unique characteristics, such as distinct responses to immunotherapies and gene expression profiles.

Recently, progress has been made in understanding TCE. For example, the definition and identification of exhausted T cells has changed from being based on the cells’ phenotype to being classified at the transcriptional and epigenetic levels^5,6^. The ability for T cells to use identical underlying genomes to generate differentiated cells with diverse gene expression profiles has prompted interest in understanding how a common set of transcription factors drives differentiation. Furthermore, various models have been proposed^6–8^ to account for the progression of T cell states, from the acute phase of immune responses to exhaustion. Two core differentiation models have been proposed to reflect the transcriptional profiles of T cell subsets: the “linear” and “circular” models^6,8^. The circular model (Figure 1A), proposes that naïve T cells (T_N_) can cycle between memory (T_M_) and exhausted (T_E_) intermediary states, resulting in an oscillating (On-Off-On or Off-On-Off) pattern of transcriptional changes over time before reaching the terminal state (T_T_). This sets up a recurring cycle of T cell differentiation (T_N_ → T_M_ ↔T_E_ →T_T_). Conversely, the linear model (Figure 1B), proposes that signal strength and duration of signals are key determining factors of T cell differentiation. This results in gene expression patterns that change gradually (On-Off, Off-On) as cells progress towards their terminally differentiated state. Moreover, this sets up the progressive loss of memory-associated gene expression and gain of exhausted-associated gene function during CD8+ T cell differentiation (T_N_ → T_M_ → T_E_ →T_T_). In both the circular and linear differentiation models, antigen-specific signals transduced through the T cell receptor (TCR) play an important role in driving the transcriptional changes that underlie exhausted T cell characteristics^9,10^. Many of the primary transcription factors involved in CD8+ T cell activation and differentiation have been identified, including TCR, activator protein 1 (AP1), nuclear factor of activated T cells 1 (NFATC1), and PD1. However, efforts to target CD8+ T cells therapeutically and inhibit their exhaustion are hindered by the lack of a mechanistic understanding of the gene regulatory pathways and differentiation subsets driving T cell exhaustion.

**Figure 1:**
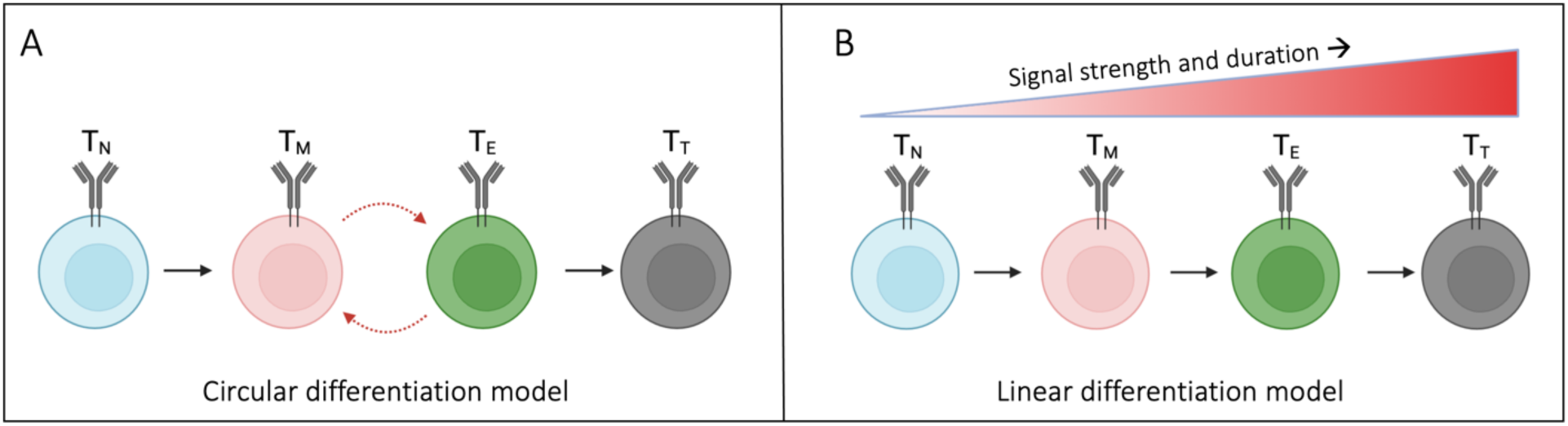
Proposed CD8+ T cell differentiation models result in unique gene expression patterns over time. (A) In the circular model of CD8+ T cell differentiation, naïve T cells (T_N_) cycle between memory (T_M_) and exhausted (T_E_) intermediates before reaching a terminal differentiated state (T_T_). (B) In the linear model, CD8+ T cells differentiate depending on the gradual acquisition of memory- or exhausted-associated genes.

Mathematical modeling is needed to understand the complex interplay of the many genes that mediate T cell exhaustion. Specifically, Boolean models have emerged as a modeling tool to study biological systems where it is of interest to understand the dynamics of cellular states and functions under different conditions. Previous Boolean models of T cell activation have been developed, integrating pathways from TCR stimulation and co-stimulatory molecules^11,12^. These models have been applied to study activation of T cells via TCR and CD28^12^ when they encounter tumor cells, and natural killer cells with CD4/CD8 co-receptors^11^. The dynamics of Boolean models are typically examined once they reach a stable behavior. At this point, the sets of interactions have converged to an attractor end state, which is interpreted as a physiological endpoint. However, it is important to also examine the transitions of cell states that lead to their attractors in order to investigate the progression of T cell states. We aim to leverage previous modeling work to systematically evaluate the transitions between transcriptional profiles of different subsets of T cells and, ultimately, gain insight into strategies to inhibit T cell exhaustion.

In this work, we adapted a literature derived, data-driven Boolean model^13^ of CD8+ T cell gene regulation stimulated by positive and inhibitory receptors, TCR and PD1 respectively. This model has not previously been applied to study the dynamics of genes that characterize TCE. We hypothesized that diverse gene regulatory interactions are responsible for TCE rather than the dysregulation from a single gene through distinct intermediary T cell states. We applied the model to quantitively investigate the evolution of T cell states of gene regulation along the path to exhaustion. Our systems analysis identifies eight attractor states comprised of distinct patterns of gene expression corresponding to T cell exhaustion, in response to T cell activation. The sequence of changes in gene expression that gives rise to these eight cell states include four paths represented by the circular model and four paths represented by the linear differentiation model. Furthermore, the attractor states are predominately characterized by activation of exhausted-associated genes. With the predictions of the transcriptional patterns leading to T cell exhaustion, we simulated interventions that inhibit T cell exhaustion. *In silico* PD1 blockade through repression of NFATC1 is predicted to lead to seven attractor end states, predominately produced by the circular differentiation model leading to terminal pro-memory attractor states. Overall, our results show that a common gene regulatory pathway model of C8+ T cell activation can display a wide array of observed behaviors seen experimentally, including therapeutically targeting T cells through PD1. As such, the model can be used to identify novel therapeutic targets.

## RESULTS

### Population level dynamics displays fractional activation of primarily exhausted-associated genes

The dynamics of the adapted Boolean model of gene regulatory interactions provide an avenue for exploring the evolution between naïve and terminally exhausted T cell states. We utilized this model to investigate transcriptional regulation of CD8+ T cell differentiation that exists through diverse gene patterns over time and to generate novel hypotheses about the gene expression changes underlying TCE. We generated time courses by simulating the model 10,000 times and examined the influence of proliferative pro-memory” (PP) and “effector exhausted” (EE) associated genes over time. Each end state (ES) is classified as either PP or EE, based on the total number of associated genes on at the end of each time course (Materials and Methods). The model predicts that in response to initial stimulation by TCR and the IL family^13^, the fractional activation of EE genes primarily dominates over time (Figure 2A). The fractional activation of EE genes exhibits a rapid-onset, short-term peak in the first four time steps following initial stimulation. In comparison, the model predicts that the fraction of PP genes on only exceeds the fraction of EE genes that are on for time steps five through seven. This suggests that the initial influence of EE-associated genes is driving the general dynamics of the T cell population. After those initial times, the overall trends for both PP and EE activation is comparable, and we see by time step 13, the fraction of genes that are active in both groups is below 10% for the remaining time steps.

**Figure 2:**
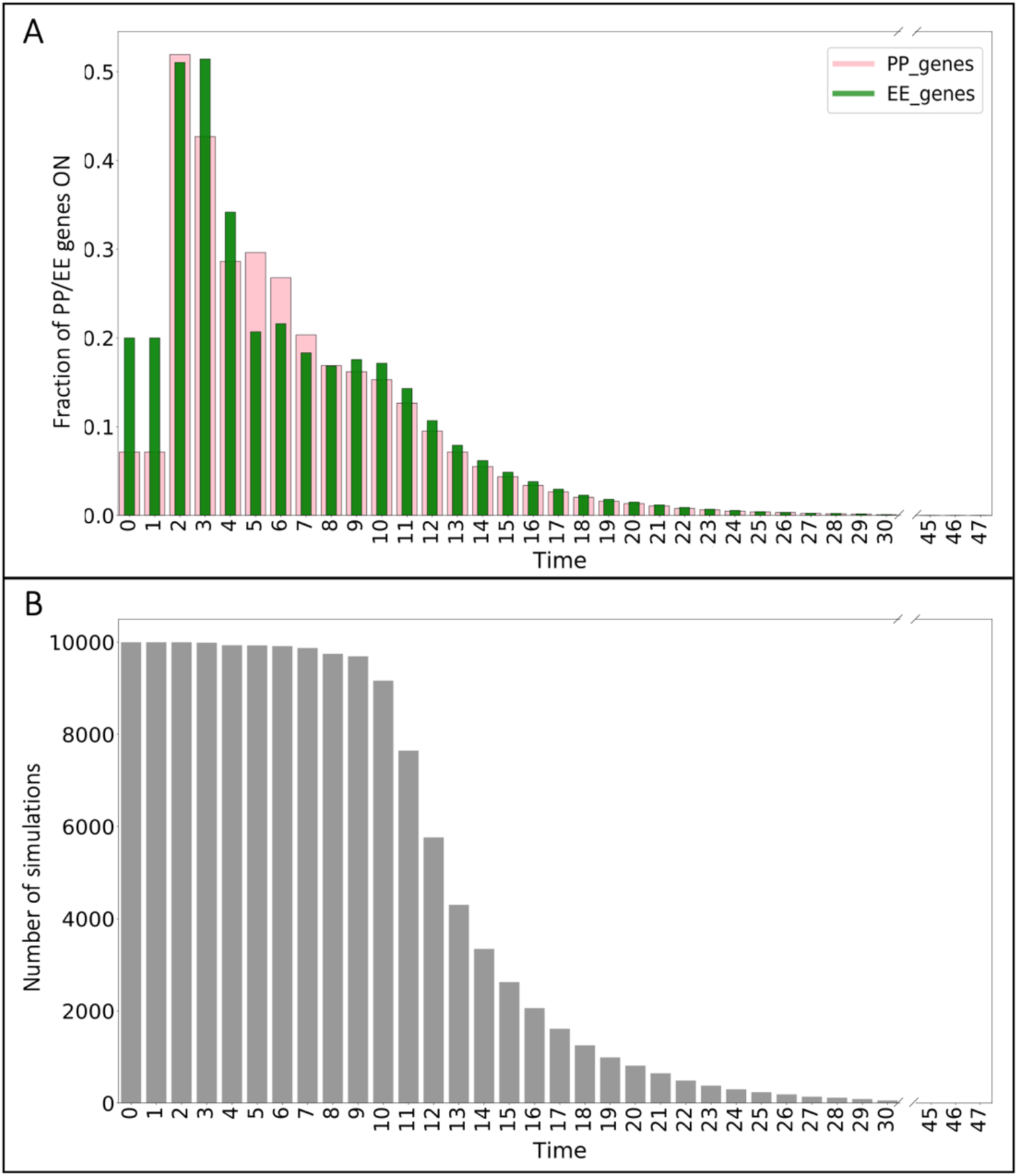
Fractional activation profiles following initial T cell activation. (A) Fraction of PP and EE gene activation for all simulations over time. (B) Number of simulations that have not reached their terminal end state.

Next, we obtained the frequency of simulations remaining at each time step (Figure 2B). This frequency represents the *in silico* cells that have not reached their terminal end state at a particular time. Notably, only a small fraction of the total simulation of cells (< 500 out of 10,000) terminate in the first nine time steps. Following this, there is a quick decline in the number of cells that have reached their end state from the initial 10,000 to less than half of the total by time step 13. At that point, only a small number of cells (< 20) remain before reaching their end state by the end of the simulated time. The results in Figure 2 show there are more EE genes turned on (> 50%) in the cells that reach a terminal end state in the first few time steps. In all, the results from examining the overall dynamics of the population reveal dominance towards T cell exhaustion from early time steps before the majority of cells reach terminal differentiation.

### Network analysis identifies eight distinct end state attractors of T cell exhaustion

Once we established the overall population-level dynamics predicted by our model, we proceeded to evaluate how the model input impacts the terminal end state for each simulated T cell. This allowed us to identify the network of interactions that comprise the final state for an individual T cell following initial stimulation. Utilizing the generated time-courses representing single cells, we aimed to determine the influence of transcriptional activation of PP and EE genes. The predicted outputs from 10,000 simulations resulted in eight attractor end states, consisting of a broad range for the number of cells in each end state (Figure 3A).

**Figure 3:**
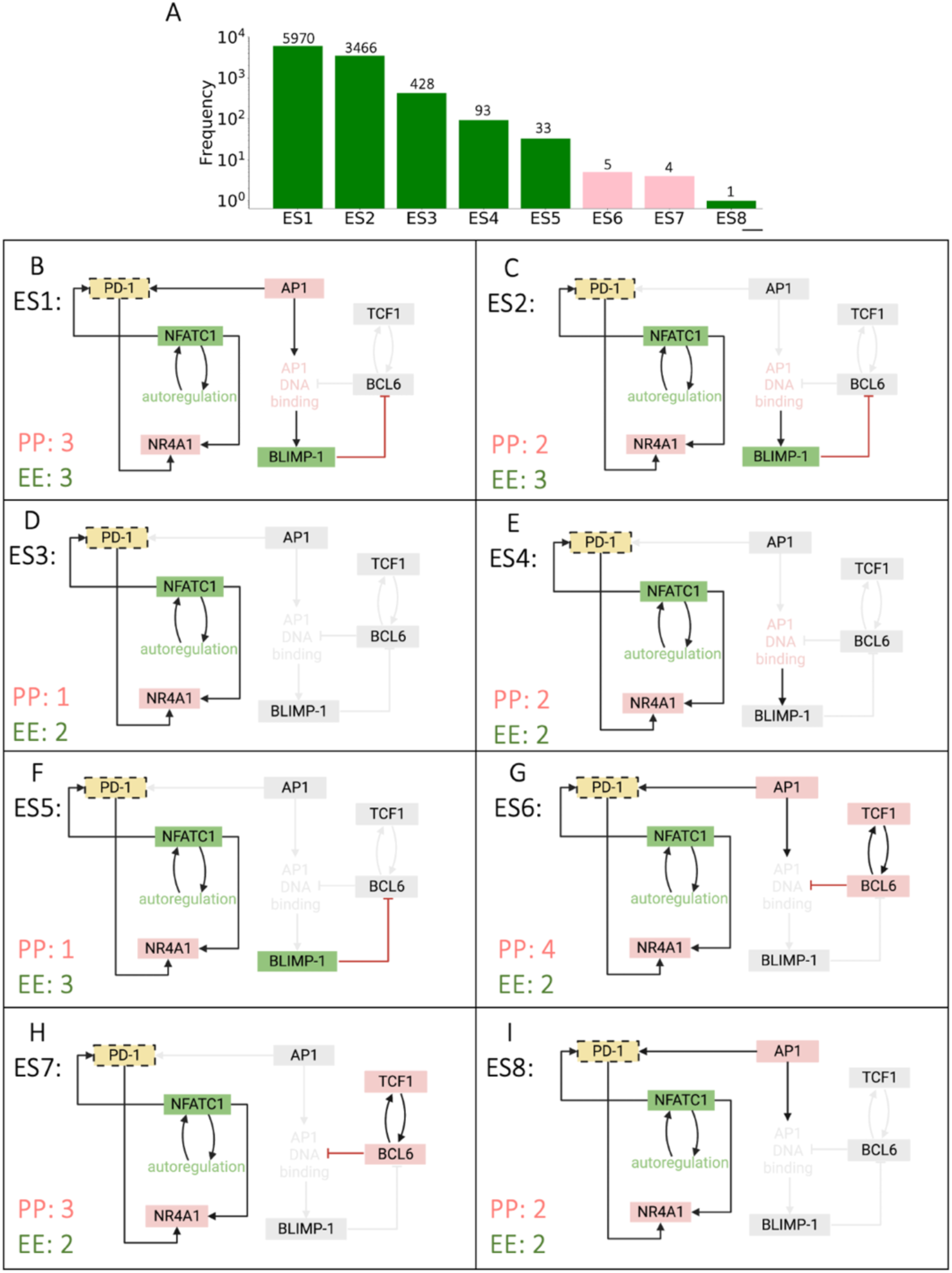
Terminal end states from Boolean model. (A) Frequency distributions of simulations residing in each end state attractor. Bars are color coded to reflect the terminally differentiated T cell state, pro-memory (*pink*), or exhausted (*green*). (B-I) Terminal end states 1-8, including the number of PP and EE genes remaining activated in each end state.

The model predicts that distinct combinations of PP and EE genes are on for each end state. Specifically, the eight end states identified by the model are defined by PD1, NFATC1, NR4A1, AP1, BLIMP1, TCF1 and BCL6 expression (Figure 3B-I). We observed three common gene modulators of terminal T cell differentiation: PD1, NFATC1, and NR4A1. These are not unexpected, since these genes create a positive feedforward loop with each other following TCR stimulation within the network structure. Interestingly, the model predicts that end states ES1 (Figure 3B), ES4 (Figure 3E), and ES8 (Figure 3I) exhibit an equal number of PP and EE genes turned on. However, these end states are classified as terminally exhausted due to AP1 and AP1 DNA binding activity, which are known to positively regulate T cell exhaustion^14,15^. This prediction agrees with experimental evidence showing that the ratio of NFATC1 and AP1 is important during the decision process for pro-memory versus exhaustion in CD8+ T cells^16,17^. AP1 is turned off in end state ES5 (Figure 3F); however, activation of BLIMP1 supports terminally exhausted T cells in this end state. Finally, the model predicts that only end states ES6 (Figure 3G) and ES7 (Figure 3H) are classified as PP cells. These are the only end states in which TCF1 and BCL6, genes classified as terminally pro-memory, remain on despite residual activation of PD1 and NFATC1. Activation of TCF1 and BCL6 would not be predicted solely by examining the model structure. Again, the model prediction agrees with experimental evidence, as activation along the TCF1-BCL6 axis has been shown to modulate T cell development and exert positive effects on differentiation to drive and maintain immune response of CD8+ T cells^18,19^. Altogether, our modeling results clearly illustrate that a common set of network interactions can exhibit significant differences in the transcriptional profiles that influence terminal T cell differentiation.

### Evolution of CD8+ T cell states reveals circular and linear differentiation models through distinct gene expression activation patterns

Next, we explored whether the evolution of gene activation predicted by the model follows circular or linear differentiation. Thus far, we have shown that the population dynamics of T cells exhibit differences in fractional activation of both PP and EE genes (Figure 2), and the end state of individual cells have different network structures (Figure 3). Together, these results suggest that activation of specific genes over time drives T cell response, leading to the predicted end state. Therefore, to investigate the evolution of T cell progression, we examined the state changes over time for each end state identified. We were particularly interested in the expression pattern for each gene as a cell progresses through intermediary states to reach its terminal end state. To this end, we chose a representative cell from each of the eight end states and tracked the changes in expression of each gene over time. We utilized these gene expression patterns to classify the T cell state at every time point.

The model predicts distinct sequences of gene expression leading to the terminal state. The first two time steps across all cells represent the naïve state (T_N_) for a T cell, following the initial stimulation by TCR, and IL family. We can then visually observe differences between the fractional activation of PP and EE genes and patterns of genes that turn on and off in each cell over time. In particular, we observed five groups of single cells (SC) (SC1 (Figure 4A), SC2 (Figure 4B), SC3 (Figure 4C), SC4 (Figure 4D), and SC5 (Figure 4E)) characterized by an oscillatory pattern of expression of PP and EE genes. These groups are classified as following the circular model of differentiation. In comparison, three other groups exhibit a more gradual increase in their expression of PP or EE genes (SC6 (Figure 4F), SC7 (Figure 4G), and SC8 (Figure 4H)), following the linear model of differentiation. Within the circular models, T cell state progression is driven by the activation of pro-memory genes, such as BCL6, TCF1, and FOXO1, as well as exhausted genes, including BATF, IRF4. This results in a cycle between the PP and EE states (T_M_ and T_E_, respectively) before reaching the terminal (T_T_) state of T cell exhaustion (*green*) state. Interestingly, the model simulations predict that the groups that follow the linear model can terminate in different end states. Specifically, the SC3 (Figure 4C) and SC8 (Figure 4H) groups terminate in an exhausted end state through residual activation of PD1 and NFATC1, while the SC6 and SC7 groups terminate in a pro-memory state (*pink*) driven by activation of TCF1 and BCL6. It is important to note that despite the equal number of PP and EE genes on in SC1, SC4, SC8 (Figure 4A,D,H), these cells are classified as terminally exhausted because of the sustained activation of AP1, a positive regulator of TCE^14^. Finally, model simulations predict that SC2 (Figure 4B), SC4 (Figure 4D), and SC5 (Figure 4E) are the only groups where AKT, mTOR, and glycolysis are active together. Intuitively, we can understand this sequence of interactions following TCR stimulation where AKT becomes active if PD1 is not active. Active AKT then triggers the activation of mTOR, which results in glycolysis. These signals immediately become deactivated following PD1 activation. Our predictions agree with experimental evidence, where studies have shown that activation of naïve CD8+ T cells can also trigger alterations in metabolism.

**Figure 4:**
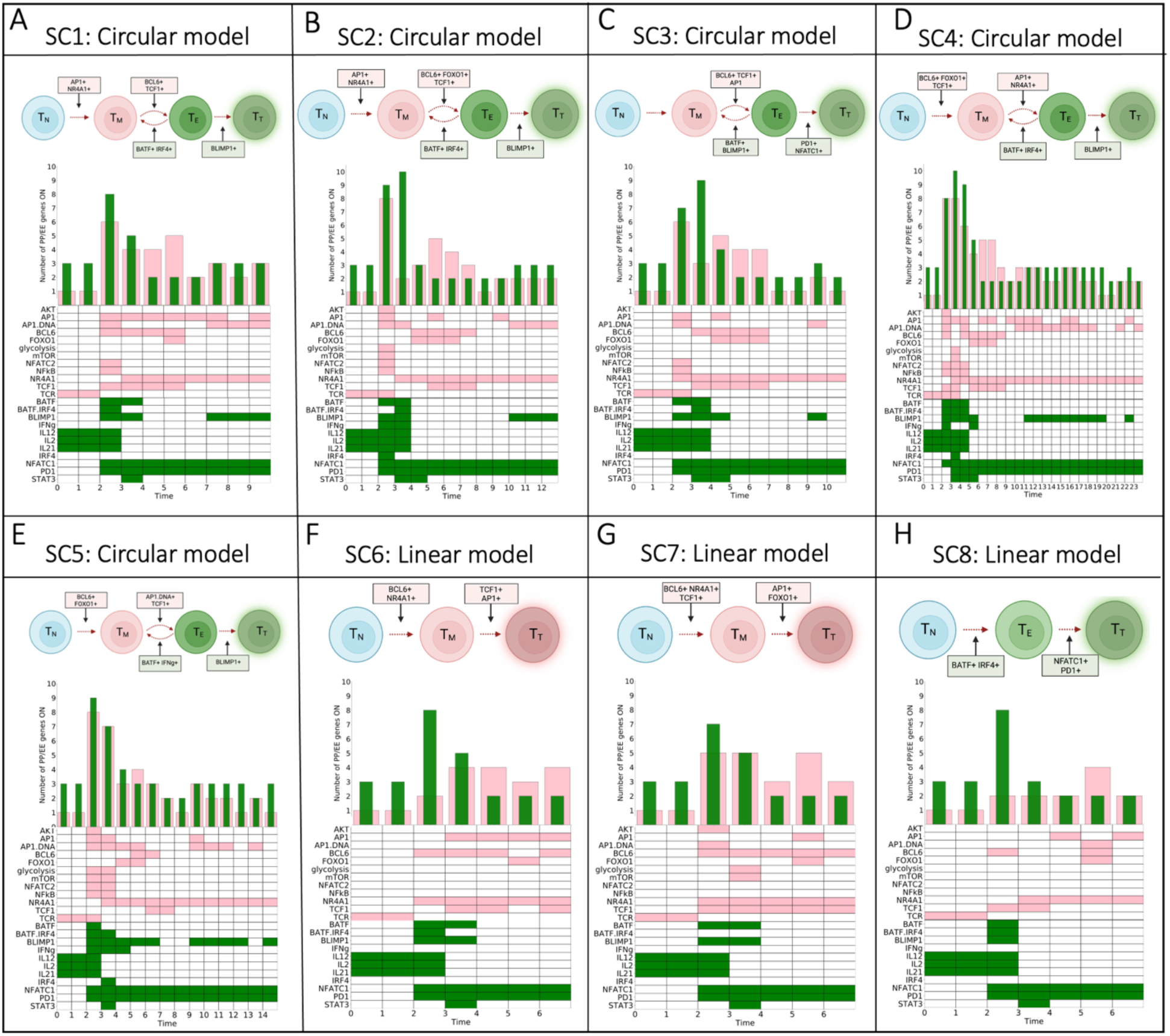
Evolution of T cell states and associated differentiation models. (A-H) Single T cell tracking intermediary states defined by PP (*pink*) or EE (*green*) gene activation. Schematic diagram represents the overall trend for each single cell time course. Frequency distribution bars and heatmap are color coded to reflect the terminally differentiated T cell state, pro-memory (T_T_, *pink*), or exhausted (T_T_, *green*).

### Population level T cell dynamics from PD1 blockade display fractional activation of memory-associated genes

Following the identification of distinct network interactions comprising two T cell differentiation models, we proceeded to evaluate how perturbing the network (Materials and Methods) impacts the dynamics of the population compared to cells with the baseline network (*wildtype*, WT). Drawing on published work, we specifically focused on perturbing interactions that affect PD1. Recent experimental studies^20^ have focused on attempting to block the PD1 pathway using anti-PD1 antibodies during the naïve-to-memory CD8+ T cell transition, based on results showing that PD1 can be modulated between these stages of T cell differentiation^21,22^. However, the role of PD1 in CD8+ T cell differentiation remains poorly characterized. To address this gap, we encoded a PD1 checkpoint blockade in the model by repressing NFATC1, a known activator of PD1^23^. We simulated time-courses similar to WT to examine the effect on transcriptional activation of PP and EE genes. The model predicts that with PD1 blockade, in response to initial stimuli, the fractional activation of PP genes primarily dominates over time (Figure 5A). The fractional activation of PP genes exhibits a quick, transient peak in the first three time steps following initial stimulation. In comparison, we observed that the fraction of activated EE genes is only prominent in time step four. This initially suggests that the influence of memory-associated genes is driving the general dynamics of the population. The long-term behavior for activation of both PP and EE genes over time is similar, and by time step 15, the fraction of active genes in both groups is below 10% for the remaining time steps.

**Figure 5:**
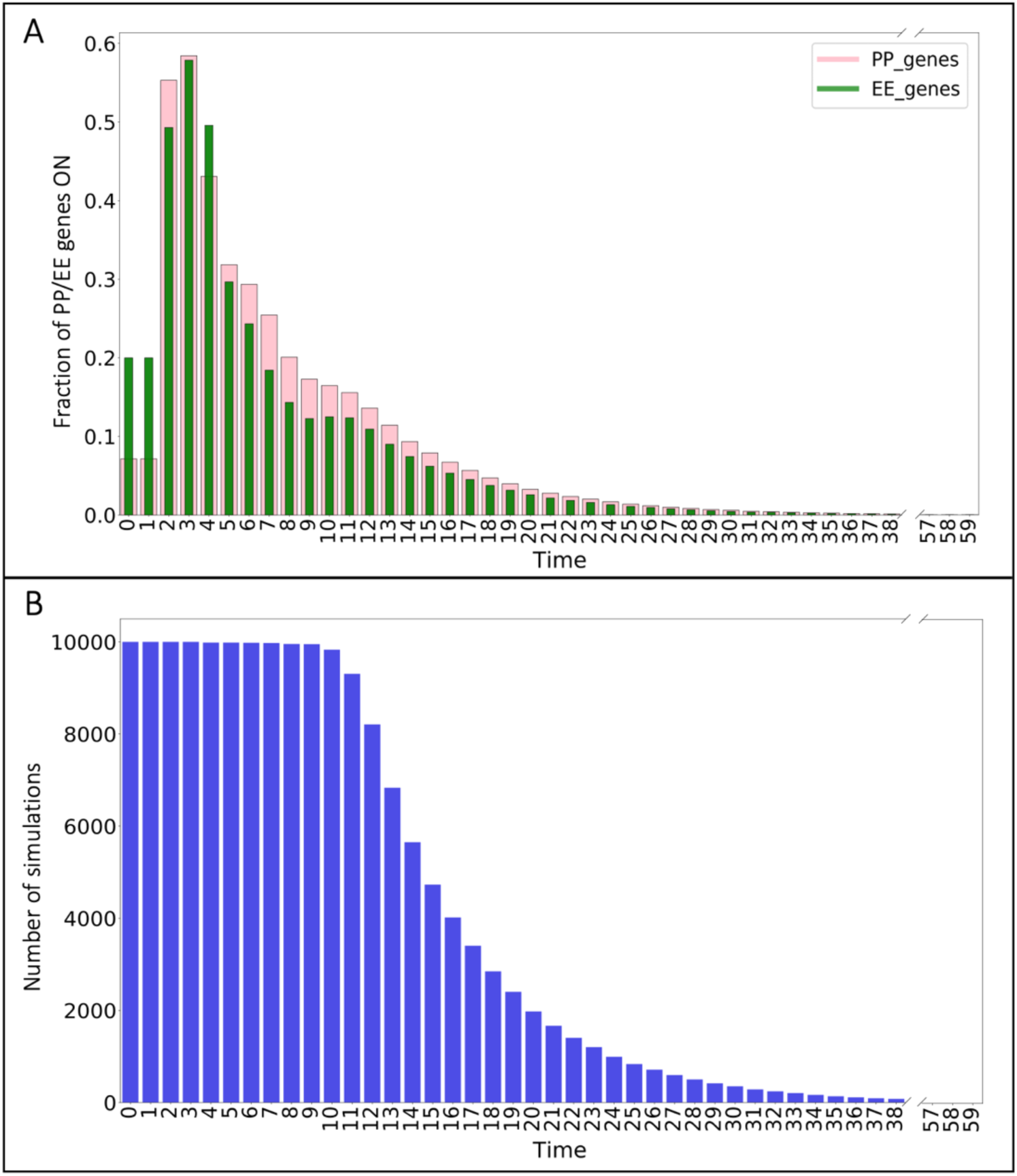
Fractional activation profiles following initial T cell activation and PD1 blockade. (A) Fraction of PP and EE gene activation for all simulations over time in response to PD1 blockade. (B) Number of simulations remaining before reaching terminal end state.

In addition to examining the influence of the PD1 blockade on the fractional activation profiles for PP and EE genes, we also observed the frequency of simulations remaining before reaching their terminal end state (Figure 5B). The results show that blocking PD1 delays cells reaching their terminal end state compared to WT (Figure 2B). Notably, we only observe a small number of the total simulations (<10) that concludes in the first 10 time steps, in contrast to the quick decline in the number of cells from the total 10,000 to less than half of the total by time step 15. It is interesting to note that more than half of the total population reaches its terminal state within the first several time steps, and only a small number of cells (< 50) remain before reaching their end state by the end of the time course. Overall, the results from examining the dynamics of the population with PD1 blockade reveal dominance of the PP genes before cells reach their end states.

### Network analysis with PD1 blockade identifies seven distinct end state attractors of T cell exhaustion

In order to elucidate the mechanisms by which T cells respond to a PD1 checkpoint blockade, we simulated a population of 10,000 single cells and examined the terminal differentiated end states. Model prediction outputs revealed seven attractor end states, with a broad frequency distribution (Figure 6A). The majority of simulation outputs (∼86%) go to end states ES1 and ES2, both terminally PP end states. We can see that the overall end state network structure has differential activation combinations of PD1, NFATC1, NR4A1, AP1, BLIMP1, BCL6 and TCF1. There are two common gene modulators across the seven end states, NFATC1 and NR4A1 (Figure 6B-I). These are interesting, because previous studies have proposed that the NFATC1-NR4A1 axis controls the T cell exhaustion program^24^ and NR4A1 has stronger control over the regulation of T cell dysfunction^25^. Therefore, we classify ES4 (Figure 6F), the only end state composed solely of the two common gene modulators, as terminally pro-memory. More intriguing is that despite the PD1 blockade, two of the end states, ES3 (Figure 6D) and ES6 (Figure 6H), remain terminally exhausted. For ES3, although we removed the ability for NFATC1 to activate PD1, AP1 can still activate PD1, which provides an alternate route for PD1 to remain active. In the case of ES6, residual activation of BLIMP1 causes this end state to remain terminally exhausted despite the absence of PD1. Interestingly, ES1 (Figure 6B) and ES2 (Figure 6C) are terminally PP end states and display similar networks, with BLIMP1 remaining activated, inhibiting the activation of pro-memory associated genes, TCF1 and BCL6. Lastly, ES7 (Figure 6I) is the only terminally PP end state with activated TCF1. In all, these results reveal that PD1 blockade influences T cell terminal end states to be classified as PP cells.

**Figure 6:**
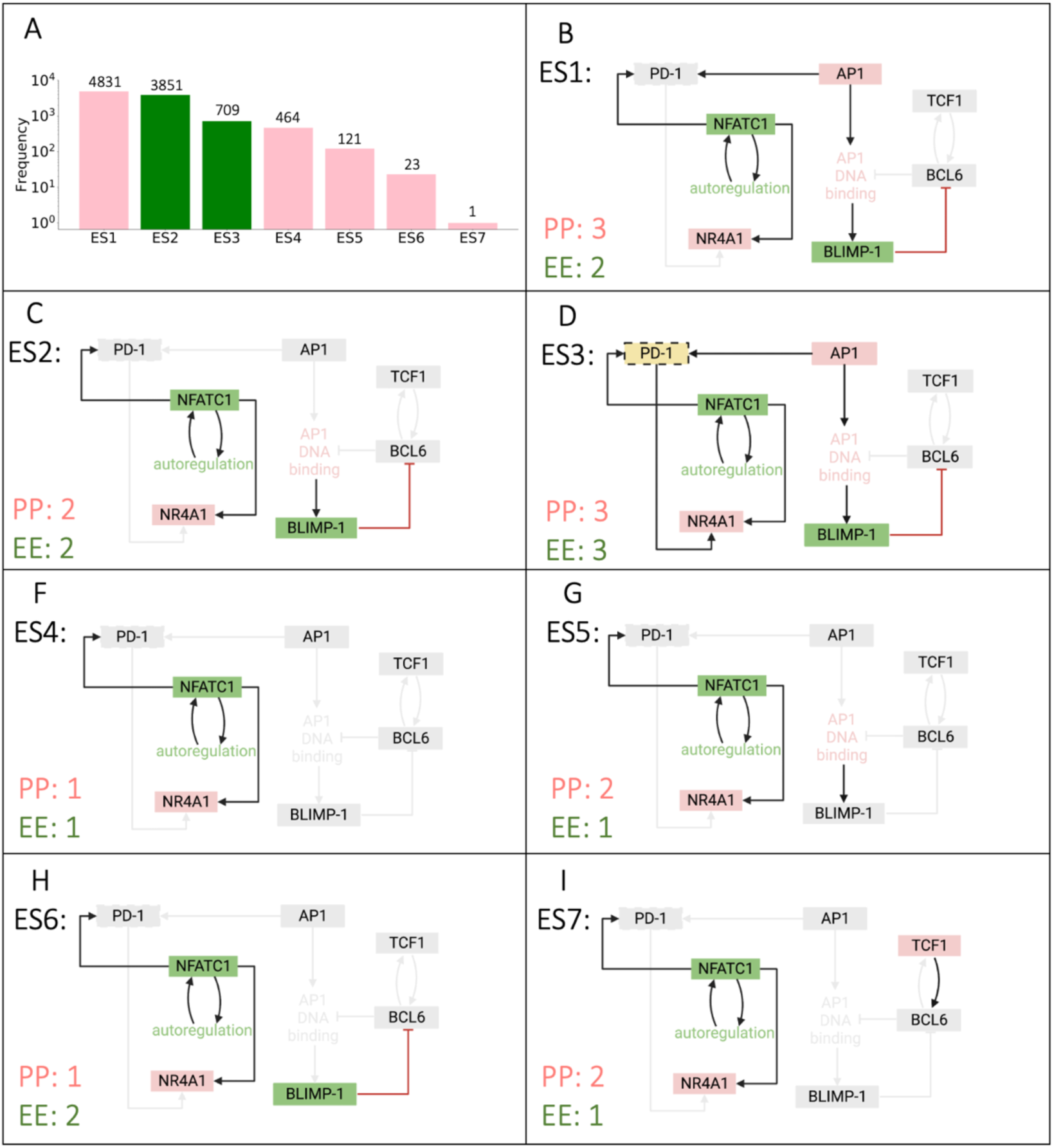
Terminal end states from Boolean model with PD1 blockade. (A) Frequency distributions of simulations residing in each end state attractor. Bars are color coded to reflect the terminally differentiated T cell state, pro-memory (*pink*), or exhausted (*green*). (B-H) Terminal end states (ES1-ES7), including the number of PP and EE genes remaining activated in each end state.

### Evolution of CD8+ T cell states with PD1 blockade reveals circular subset differentiation models

We have shown that the population dynamics of T cell response to a PD1 blockade exhibit dominance of PP gene activation (Figure 5A) and altered end state networks compared to wildtype (Figure 6). An interesting and unexplored aspect of targeting PD1 in T cell biology is that this inhibitory receptor is expressed not only on chronically stimulated exhausted CD8+ T cells, but also during the early stages of T cell activation^21,26^. One of the main goals of therapeutically targeting PD1 is to understand whether blocking PD1 during the early phase has an inhibitory role similar to what is seen in exhausted CD8+ T cells. To address these questions, we investigated the evolution of T cell response with a PD1 blockade. We sought to investigate the hypothesis that if PD1 was blocked through NFATC1 repression, the intermediary states and terminal end state of these CD8 + T cells would shift from EE to a terminal PP state. We were particularly interested to see which genes become activated as T cell progress through their intermediary states. Towards this end, we chose a representative single cell from each end state group (Figure 7A) to examine the overall progression of T cell states. Similar to WT cells, we were able to utilize these gene expression patterns to classify the state of each group of T cell end states, for every time point.

**Figure 7:**
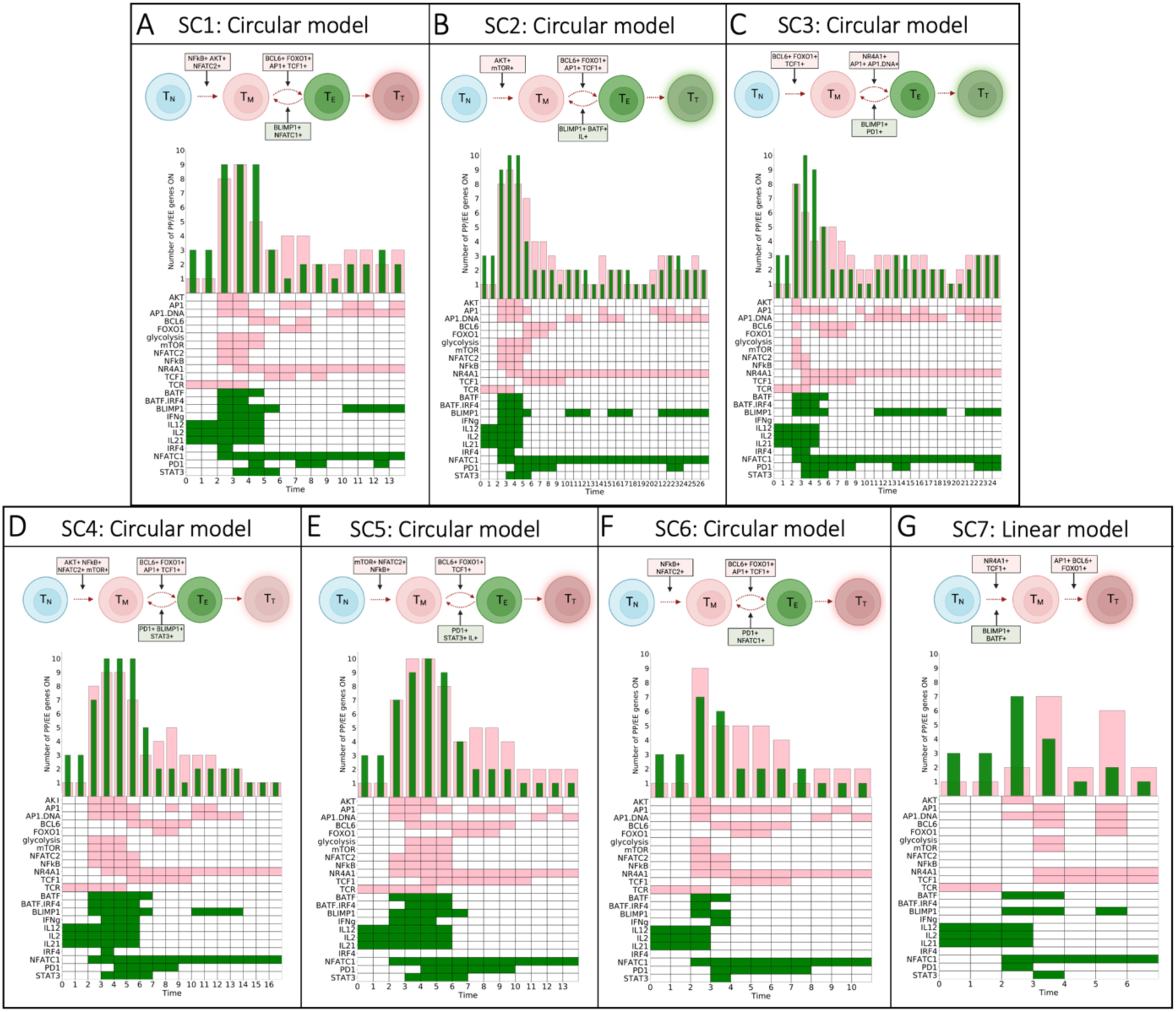
Evolution of T cell states and associated differentiation memory models with PD1 block. (A-H) Single cell tracking of intermediary T cell states defined by PP (*pink*) or EE (*green*) gene activation with PD1 blockade. Schematic diagram represents the overall trend for each single cell time course and corresponding differentiation model. Frequency distribution bars and heatmap are color coded to reflect the terminally differentiated T cell state, pro-memory (T_T_, *pink*), or exhausted (T_T_, *green*).

We can visually observe differences between the activation patterns of PP and EE genes in each group over time predominantly displaying pro-memory activation (Figure 7). In particular, we observed six of the seven groups follow the circular model, SC1 through SC6 (Figure 7A-F), while SC7 (Figure 7G) follows the linear model. Within the circular models, we observed increased activity of metabolic genes in early time steps (NFkB, AKT, mTOR, and glycolysis) before cycling between PP and EE genes. This indicated that the PD1 blockade allowed for prolonged activation of TCR and downstream activation of metabolism-associated genes compared to WT cells. Cycling between PP and EE T cell states was primarily driven by PP genes, BCL6, FOXO1, TCF1 and EE genes, BATF1, BLIMP1, NFATC1, and STAT3 (Figure 7A-F). This resulted in a cycle between T_M_ ↔ T_E_ before reaching the terminally PP (*pink*) state. Furthermore, the representative single cell that followed the linear model, SC7 (Figure 7G), terminates in a PP end state through residual activation of TCF1 and NR4A1. Lastly, it has been shown that PD1 is required for optimal CD8+ T cell memory^21^. Interestingly, we observed oscillatory dynamics of PD1 in each group, with differing activation time and length. We can understand this intuitively through AP1 activity, where PD1 turns on and off following AP1 activation and inactivation. In summary, we predict that for all single cell groups, PD1 is required to reach a pro-memory terminal end state, as T cells evolve through intermediary states.

## DISCUSSION

We applied a computational modeling approach to explore T cell states, motivated by published experimental studies. A recent review of CD8+ T cell memory^27^ described how CD8+ T cells containing identical underlying genomes can follow different patterns of differentiation, ultimately generating subsets of cells characterized by distinct gene expression. Importantly, the authors emphasized that regulation mechanisms underlying these subsets, which include both memory and exhausted T cells, are incompletely understood. However, they proposed that changes in transcription factors drive the functional differentiation and heterogeneity of these T cell subsets. The results presented in this work are consistent with this view: a data-driven Boolean model comprising core CD8+ T cell genes expressed in response to stimulation (Figure 8), predicts the fractional activation of memory and exhausted-associated genes (Figure 1), including differentiation models (Figure 3) that lead to terminal states (Figures 2 and 4). Furthermore, the model provides a framework for predicting terminal end states and differentiation dynamics of cells with altered gene regulatory networks (Figures 5 and 7). Thus, the dynamics predicted by the model align with experimental observations and can be used to investigate the evolution of T cell activation and exhaustion.

**Figure 8:**
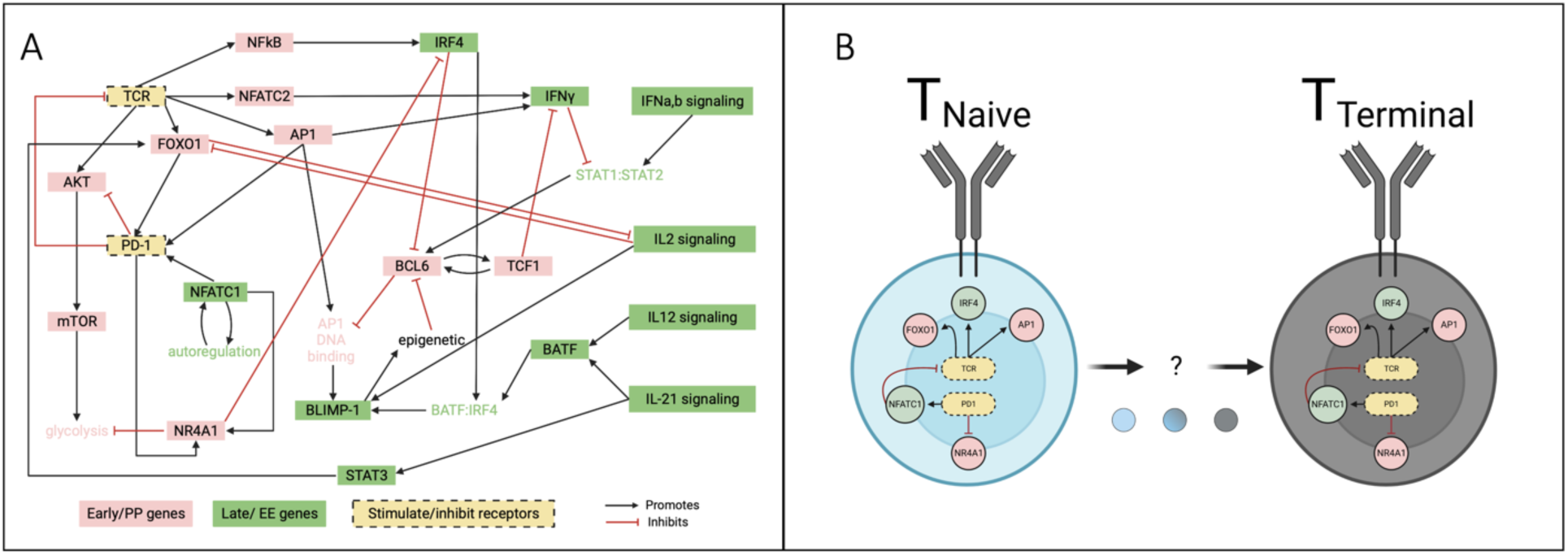
Boolean model of CD8+ T cell exhaustion. (A) Literature-based network diagram of key gene regulatory interactions underlying TCE. Black arrows indicate activation, red lines with bars on the end indicate inhibition. Stimulating and inhibitory immune receptors are shown in yellow with black dashed borders. Pink nodes represent proliferative pro-memory associated genes, green nodes represent effector exhausted associated genes. (B) Schematic outline of T cell state progression from naïve to terminal differentiation.

In this study, we adapted a data-driven Boolean model of CD8+ T cells^13^ to examine transcriptional gene patterns and differentiation along the path from naïve to exhaustion. We applied the model to investigate various characteristics of T cell signaling. By examining the transcriptional gene pattern activation of PP- and EE-related genes, we found the population of *in silico* T cells associated with eight distinct terminal gene networks (Figure 3), even when starting from identical underlying networks. These networks comprise global gene modulators, including known effectors such as PD1, NFATC1, and NR4A1. Particularly, our results highlight NFATC1 as a regulatory element that can be targeted, as it is responsible for increasing transcription of NR4A1 and regulating PD1 gene expression. This is consistent with previous transcriptomics studies^25^ that have suggested that NR4A1 may have an important role in the early stages of the CD8+ T cell response through cooperation with NFATC1, contributing to the dysfunctional state of T cells.

The PD1 pathway is known to regulate dysfunctional T cells in chronic infection and cancer, but the role of this pathway during acute infection remains less clear. PD1 is expressed by all T cells during activation, positioning this pathway to play an essential role in shaping fundamental properties of CD8+ T cell responses early during differentiation. Previous work^21^ showed that early T cell responses in absence of PD1 signals, result in defects in CD8+ T cell memory; however, a PD1 blockade only during the early stages of infection resulted in optimal memory. PD1 gene regulation has been shown in part to occur via the recruitment of NFATC1 to provide the molecular mechanism responsible for the induction of PD1 in response to T cell stimulation^23^. Our modeling analysis with a PD1 blockade through NFATC1 repression during early T cell response results in a terminal pro-memory differentiated state (Figure 7), consistent with our initial hypothesis. The activation of PD1 reveals an oscillatory pattern over time, resulting in a terminal pro-memory end state for all seven subsets. This suggests that the PD1 pathway may continue to function as a link to signals required for memory T cell maintenance, consistent with previous experimental observations^21,22,28^. In addition, early T cell response results in increased metabolic activity prior to PD1 activation. Naïve T cells have been shown to possess a metabolically resting phenotype, activated upon TCR stimulation^29^. This implies that our model analysis supports the notion that metabolism is a strong driver of CD8+ T cell differentiation and function. Lastly, it is interesting to note that we observed an even distribution of linear and circular differentiation models in WT cells; however, with PD1 blockade, the differentiation path shifts towards predominately circular. We attribute the shift to the oscillatory dynamics of PD1 activation and inactivation when PD1 is repressed by NFATC1. A future study could investigate the T cell response at different time steps clarifying the mechanisms by which NFATC1 repression of PD1 affects the T_N_ → T_M_ → T_E_ stages of CD8^+^ T cell differentiation and impact on metabolic activity. This will be important for determining how to optimally administer PD1 blocking agents alone and in combination with other therapies to achieve durable improved immunity.

We examined how the gene expression pattern changes between the intermediary T cell states along the path to exhaustion. Our modeling analysis emphasizes the influence of AP1 and BLIMP1 gene activation in driving terminal exhaustion, despite opposition with activation of pro-memory associated genes, TCF1 and BCL6. It is important to note that the timing of BLIMP1 versus TCF1 and BCL6 activation is crucial in determining the progression of terminal T cell differentiation. Particularly, in SC7 (Figure 4G) and SC8 (Figure 4H), we observed BLIMP1 activation and deactivation only during early stages of T cell response, with subsequent activation of TCF1 and BCL6. Previous studies^30^ have shown that the TCF1-BCL6 axis can repress T cell exhaustion and maintain T cell immune response. These results indicate that targeting this axis is a mechanism by which T cell-mediated immunity may be enhanced during chronic infections and cancer. Thus, we provide quantitative insights to motivate directly targeting TCF1 and BCL6 in combination with BLIMP1 to further investigate these genes as therapeutic targets.

Along with the significant findings produced by our work, we recognize some aspects that can be addressed in the future. For example, the proposed Boolean model of CD8+ T cells can be expanded to include additional genes and signaling pathways known to play a role in T cell activation and response^31–34^ (i.e., Notch, and MAPK), and additional T cell ligands^10^ (i.e., MHC class I, and MCH class II) and receptors^35–38^ (i.e., CD4, CTL4A, and CD45). Additionally, a limitation in our study is our definition of pro-memory or exhausted T cell state defined solely based on the relative number of PP and EE genes activated. Our motivation for determining T cell state based on PP and EE gene count is the fact that gene expression is the process by which information held a gene is converted into a function. Thus, we reason that the gene activation count can be determinant of T cell state. Future work can explore the possibility of a weighted probability of cell state transition^39^ in T cells, or incorporate a steady-state probability distribution function^39^ to estimate the transition potential for each T cell state in the Boolean network model.

Progress in T cell immunology research in the past decade has led to significant advancement in our understanding about various molecular factors driving T cell activation and memory formation. Despite the markedly high response rates and improved overall patient survival following checkpoint blockade, most patients did not experience durable cancer regression, and blocking PD1 alone did not overcome CD8^+^ T cells exhaustion^20,28,40^. The plasticity of CD8+ T cells strongly encourages alternate strategies for targeting CD8+ T cell subsets for future immunotherapy treatments. Improving our understanding of the gene regulatory pathways that drive CD8+ T cell activation and exhaustion and other T cell signaling pathways is thus critical for developing improved therapies against these deadly diseases. We have shown that a consensus model of CD8+ T cell gene activation can display a variety of behaviors seen experimentally and can predict novel targets for modulating CD8+ T cell exhaustion. Overall, our work provides a solid foundation for future work to study the regulatory pathways that drive T cell decisions and response.

## MATERIALS AND METHODS

### Boolean model of CD8+ T cell exhaustion

We used a large-scale network model of gene regulation to investigate the progression of T cell states and understand the evolution of gene expression patterns that lead to terminally exhausted T cells. We adapted a previously published Boolean model^13^, which was constructed through an extensive literature-based network of gene regulatory interactions underpinning TCE in CD8+ T cells (Figure 8A,B). Each gene node in the network is color-coded based on its impact along the path to TCE, labeled as either “proliferative pro-memory” (PP, *pink*) or “effector exhausted” (EE, *green*). This 25-node network integrates immune inputs (TCR and PD1, *yellow*) with downstream signaling events involved in T cell activation and exhaustion. The model incorporates signaling events stimulated by the TCR, including activation of key pro-memory genes, FOXO1^41–43^, AKT^44,45^, AP1^14,16^, and NFATC2^46,47^. Subsequent activation of these genes results in mTOR^48^ activation leading to increased glycolysis^49^ or self-renewed memory through BCL6^19,50^ and TCF1^18,51,52^. In contrast, stimulation by PD1^26,53^ or the IL signaling family, IL2^54,55^, IL12^56,57^, IL21^58,59^, promotes activation of well-known drivers of T cell exhaustion including NFATC1^23,47^, BLIMP1^60^, BATF^61,62^, and IRF4^63,64^.

### Boolean model simulations

The variables in this model can take two values: 0 (inactive or absent) or 1 (active or present). Boolean logic simulations are designed to explore the characteristics of network attractor states, however, understanding the acute and chronic CD8+ T cell responses of interest requires examining the specific intermediate transitory states. Ten thousand model simulations were implemented as a series of logic statements that are executed in an asynchronous update paradigm, known to be representative of biological systems^65^. This assumes only one random component of the network is updated at each single time step. The update mechanism results in stochastic dynamics and leads to *n* possible attractor end states. An attractor end state is reached if the network evolves to a single stable state, known as a point attractor. We extended the implemented R^66^ code corresponding to the adapted Boolean model^13^ to enable more flexible exploration of T cell state changes. The WT and perturbed model terminate at the maximum 100 time steps if an attractor state is not reached before the end of the time course.

### T cell state differentiation network analysis

To determine the gene regulatory interactions that contribute to pro-memory and exhausted CD8+ T cell states, we proceeded through the following steps. First, the ensemble of Boolean model simulations from R, containing the time courses of gene expression patterns, were exported for analysis using the Python package *Seaborn*^67^. We examined the trajectories for each cell transition leading to the attractor state, in order to investigate the evolution of network behavior. Each model simulation with the same attractor end state patterns were grouped together, resulting in eight end states for WT, and seven end states for the *in silico* PD1 blockade. Next, for each group, we calculated the number of PP and EE genes that were on for each simulation and every time step. The dominating number of genes that were activated at each time step determined the intermediate states of each T cell before reaching the terminal differentiated state. PP and EE gene labeling has been previously published^13^, supporting our approach to assess the state of a T cell at each time step. Then, we categorized the state at each time step as PP or EE by counting the relative number of PP- and EE-associated genes. The sequence of intermediate states was used to determine the differentiation model classification. If the pattern of dominating genes leads to a progressive loss of PP- or EE-associated genes, these cells were classified as following the linear differentiation model. Conversely, if the pattern of dominating genes cycled between pro-memory and exhausted gene activation over the time course, these cells were classified into the circular differentiation model. We repeated this for all 10,000 model simulations to identify both the terminal and intermediate T cell states and classify the type of differentiation model T cells utilize to reach their terminal state.

## AUTHOR CONTRIBUTIONS

G.I. and S.D.F. conceived of the study. G.I. planned and carried out computational model simulations. S.D.F. supervised the project and provided financial support. All authors discussed the results and contributed to the final manuscript.

## ACKNOWLEDGEMENTS

The authors thank members of the Finley research group for constructive feedback and Dr. Hamid Bolouri for assistance with accessing files from the Boolean model.

## REFERENCES

1. Dolina, J. S., Van Braeckel-Budimir, N., Thomas, G. D. & Salek-Ardakani, S. CD8+ T Cell Exhaustion in Cancer. Front. Immunol. 12, (2021).

2. Philip, M. & Schietinger, A. CD8+ T cell differentiation and dysfunction in cancer. Nat. Rev. Immunol. 2021 224 22, 209–223 (2021).

3. Thommen, D. S. & Schumacher, T. N. T cell dysfunction in cancer. Cancer Cell 33, 547– 562 (2018).

4. Marin-Acevedo, J. A., Soyano, A. E., Dholaria, B., Knutson, K. L. & Lou, Y. Cancer immunotherapy beyond immune checkpoint inhibitors. J. Hematol. Oncol. 11, 8 (2018).

5. Martin, M. D. & Badovinac, V. P. Defining memory CD8 T cell. Front. Immunol. 9, (2018).

6. Henning, A. N., Roychoudhuri, R. & Restifo, N. P. Epigenetic control of CD8+ T cell differentiation. Nat. Rev. Immunol. 2018 185 18, 340–356 (2018).

7. Kaech, S. M. & Cui, W. Transcriptional control of effector and memory CD8+ T cell differentiation. Nat. Rev. Immunol. 2012 1211 12, 749–761 (2012).

8. Chen, Y., Zander, R., Khatun, A., Schauder, D. M. & Cui, W. Transcriptional and Epigenetic Regulation of Effector and Memory CD8 T Cell Differentiation. Front. Immunol. 9, (2018).

9. Adachi, K. & Davisa, M. M. T-cell receptor ligation induces distinct signaling pathways in naïve vs. antigen-experienced T cells. Proc. Natl. Acad. Sci. U. S. A. 108, 1549–1554 (2011).

10. Shah, K., Al-Haidari, A., Sun, J. & Kazi, J. U. T cell receptor (TCR) signaling in health and disease. Signal Transduct. Target. Ther. 2021 61 6, 1–26 (2021).

11. Saez-Rodriguez, J. et al. A Logical Model Provides Insights into T Cell Receptor Signaling. PLoS Comput. Biol. 3, 1580–1590 (2007).

12. Martínez-Méndez, D., Villarreal, C., Mendoza, L. & Huerta, L. An Integrative Network Modeling Approach to T CD4 Cell Activation. Front. Physiol. 11, 380 (2020).

13. Bolouri, H. et al. Integrative network modeling reveals mechanisms underlying T cell exhaustion. Sci. Reports 2020 101 10, 1–15 (2020).

14. Atsaves, V., Leventaki, V., Rassidakis, G. Z. & Claret, F. X. AP-1 Transcription Factors as Regulators of Immune Responses in Cancer. Cancers (Basel). 11, (2019).

15. Lugli, E., Galletti, G., Boi, S. K. & Youngblood, B. A. Stem, effector and hybrid states of memory CD8 + T cells. doi:10.1016/j.it.2019.11.004

16. Mognol, G. P. et al. Targeting the NFAT:AP-1 transcriptional complex on DNA with a small-molecule inhibitor. Proc. Natl. Acad. Sci. U. S. A. 116, 9959–9968 (2019).

17. Seo, W., Jerin, C. & Nishikawa, H. Transcriptional regulatory network for the establishment of CD8+ T cell exhaustion. Exp. Mol. Med. 2021 532 53, 202–209 (2021).

18. Escobar, G., Mangani, D. & Anderson, A. C. T cell factor 1 (Tcf1): a master regulator of the T cell response in disease. Sci. Immunol. 5, (2020).

19. Liu, Z. et al. Cutting Edge: Transcription Factor BCL6 Is Required for the Generation, but Not Maintenance, of Memory CD8+ T Cells in Acute Viral Infection. J. Immunol. 203, 323–327 (2019).

20. Topalian, S. L. et al. Safety, Activity, and Immune Correlates of Anti–PD-1 Antibody in Cancer. N. Engl. J. Med. 366, 2443–2454 (2012).

21. Pauken, K. E. et al. The PD-1 Pathway Regulates Development and Function of Memory CD8+ T Cells following Respiratory Viral Infection. Cell Rep. 31, 107827 (2020).

22. Xu-Monette, Z. Y., Zhang, M., Li, J. & Young, K. H. PD-1/PD-L1 blockade: Have we found the key to unleash the antitumor immune response? Front. Immunol. 8, 1597 (2017).

23. Oestreich, K. J., Yoon, H., Ahmed, R. & Boss, J. M. NFATc1 Regulates Programmed Death-1 Expression Upon T Cell Activation. J. Immunol. 181, 4832 (2008).

24. Liu, X. et al. Genome-wide analysis identifies NR4A1 as a key mediator of T cell dysfunction. Nat. 2019 5677749 567, 525–529 (2019).

25. Odagiu, L., May, J., Boulet, S., Baldwin, T. A. & Labrecque, N. Role of the Orphan Nuclear Receptor NR4A Family in T-Cell Biology. Front. Endocrinol. (Lausanne). 11, 1107 (2021).

26. Ahn, E. et al. Role of PD-1 during effector CD8 T cell differentiation. Proc. Natl. Acad. Sci. U. S. A. 115, 4749–4754 (2018).

27. Montacchiesi, G. & Pace, L. Epigenetics and CD8+ T cell memory*. Immunol. Rev. 305, 77–89 (2022).

28. Odorizzi, P. M., Pauken, K. E., Paley, M. A., Sharpe, A. & John Wherry, E. Genetic absence of PD-1 promotes accumulation of terminally differentiated exhausted CD8+ T cells. J. Exp. Med. 212, 1125–1137 (2015).

29. Reina-Campos, M., Scharping, N. E. & Goldrath, A. W. CD8+ T cell metabolism in infection and cancer. Nat. Rev. Immunol. 2021 2111 21, 718–738 (2021).

30. Wu, T. et al. The TCF1-Bcl6 axis counteracts type I interferon to repress exhaustion and maintain T cell stemness. Sci. Immunol. 1, (2016).

31. Backer, R. A. et al. A central role for Notch in effector CD8+ T cell differentiation. Nat. Immunol. 15, 1143 (2014).

32. Tsukumo, S. I. & Yasutomo, K. Regulation of CD8+ T Cells and Antitumor Immunity by Notch Signaling. Front. Immunol. 9, 1 (2018).

33. D’Souza, W. N., Chang, C.-F., Fischer, A. M., Li, M. & Hedrick, S. M. The Erk2 MAPK Regulates CD8 T Cell Proliferation and Survival. J. Immunol. 181, 7617 (2008).

34. Rohrs, J. A., Siegler, E. L., Wang, P. & Finley, S. D. ERK Activation in CAR T Cells Is Amplified by CD28-Mediated Increase in CD3ζ Phosphorylation. iScience 23, 101023 (2020).

35. Gattinoni, L. et al. CTLA-4 dysregulation of self/tumor-reactive CD8+ T-cell function is CD4+ T-cell dependent. Blood 108, 3818 (2006).

36. Ledbetter, J. A. et al. CD4, CD8 and the role of CD45 in T-cell activation. Curr. Opin. Immunol. 5, 334–340 (1993).

37. Laidlaw, B. J., Craft, J. E. & Kaech, S. M. The multifaceted role of CD4+ T cells in the regulation of CD8+ T cell memory maturation. Nat. Rev. Immunol. 16, 102 (2016).

38. Rowshanravan, B., Halliday, N. & Sansom, D. M. CTLA-4: a moving target in immunotherapy. Blood 131, 58–67 (2018).

39. Joo, J. Il, Zhou, J. X., Huang, S. & Cho, K. H. Determining Relative Dynamic Stability of Cell States Using Boolean Network Model. Sci. Reports 2018 81 8, 1–14 (2018).

40. Huang, Y., Jia, A., Wang, Y. & Liu, G. CD8+ T cell exhaustion in anti-tumour immunity: The new insights for cancer immunotherapy. Immunology 168, 30–48 (2023).

41. Michelini, R. H., Doedens, A. L., Goldrath, A. W. & Hedrick, S. M. Differentiation of CD8 memory T cells depends on Foxo1. J. Exp. Med. 210, 1189–1200 (2013).

42. Kim, M. V., Ouyang, W., Liao, W., Zhang, M. Q. & Li, M. O. The Transcription Factor Foxo1 Controls Central Memory CD8+ T Cell Responses to Infection. Immunity 39, 286– 297 (2013).

43. Staron, M. M. et al. The Transcription Factor FoxO1 Sustains Expression of the Inhibitory Receptor PD-1 and Survival of Antiviral CD8+ T Cells during Chronic Infection. Immunity 41, 802–814 (2014).

44. Rogel, A. et al. Akt signaling is critical for memory CD8+ T-cell development and tumor immune surveillance. Proc. Natl. Acad. Sci. U. S. A. 114, E1178–E1187 (2017).

45. Abdullah, L., Hills, L. B., Winter, E. B. & Huang, Y. H. Diverse Roles of Akt in T cells. Immunometabolism 3, (2021).

46. Agnellini, P. et al. Impaired NFAT nuclear translocation results in split exhaustion of virus-specific CD8+ T cell functions during chronic viral infection. Proc. Natl. Acad. Sci. USA 104, 4565–4570 (2007).

47. Peng, S. L., Gerth, A. J., Ranger, A. M. & Glimcher, L. H. NFATc1 and NFATc2 together control both T and B cell activation and differentiation. Immunity 14, 13–20 (2001).

48. Kim, E. H. & Suresh, M. Role of PI3K/Akt signaling in memory CD8 T cell differentiation. Front. Immunol. 4, 20 (2013).

49. Dang, C. V. Links between metabolism and cancer. Genes Dev. 26, 877–890 (2012).

50. Ichii, H. et al. Role for Bcl-6 in the generation and maintenance of memory CD8+ T cells. Nat. Immunol. 2002 36 3, 558–563 (2002).

51. Chen, Z. et al. TCF-1-Centered Transcriptional Network Drives an Effector versus Exhausted CD8 T Cell-Fate Decision. Immunity 51, 840–855.e5 (2019).

52. Zhang, J., Lyu, T., Cao, Y. & Feng, H. Role of TCF-1 in differentiation, exhaustion, and memory of CD8+ T cells: A review. FASEB J. 35, (2021).

53. Alsaab, H. O. et al. PD-1 and PD-L1 checkpoint signaling inhibition for cancer immunotherapy: mechanism, combinations, and clinical outcome. Front. Pharmacol. 8, (2017).

54. Liu, Y. et al. IL-2 regulates tumor-reactive CD8+ T cell exhaustion by activating the aryl hydrocarbon receptor. Nat. Immunol. 2021 223 22, 358–369 (2021).

55. Kalia, V. & Sarkar, S. Regulation of Effector and Memory CD8 T Cell Differentiation by IL-2—A Balancing Act. Front. Immunol. 9, 2987 (2018).

56. Henry, C. J., Ornelles, D. A., Mitchell, L. M., Brzoza-Lewis, K. L. & Hiltbold, E. M. IL-12 Produced by Dendritic Cells Augments CD8+ T cell Activation through the Production of the Chemokines CCL1 and CCL17. J. Immunol. 181, 8576 (2008).

57. Hombach, A. et al. IL12 integrated into the CAR exodomain converts CD8+ T cells to poly-functional NK-like cells with superior killing of antigen-loss tumors. Mol. Ther. 30, 593–605 (2022).

58. Casey, K. A. & Mescher, M. F. IL-21 Promotes Differentiation of Naive CD8 T Cells to a Unique Effector Phenotype. J. Immunol. 178, 7640–7648 (2007).

59. Tian, Y. & Zajac, A. J. IL-21 and T cell differentiation: consider the context. Trends Immunol. 37, 557 (2016).

60. Shin, H. et al. A Role for the Transcriptional Repressor Blimp-1 in CD8+ T Cell Exhaustion during Chronic Viral Infection. Immunity 31, 309–320 (2009).

61. Kurachi, M. et al. The transcription factor BATF operates as an essential differentiation checkpoint in early effector CD8+ T cells. Nat. Immunol. 2014 154 15, 373–383 (2014).

62. Tsao, H. W. et al. Batf-mediated epigenetic control of effector CD8+ T cell differentiation. Sci. Immunol. 7, (2022).

63. Huber, M. & Lohoff, M. IRF4 at the crossroads of effector T-cell fate decision. Eur. J. Immunol. 44, 1886–1895 (2014).

64. Seo, H. et al. BATF and IRF4 cooperate to counter exhaustion in tumor-infiltrating CAR T cells. Nat. Immunol. 2021 228 22, 983–995 (2021).

65. Schwab, J. D., Kühlwein, S. D., Ikonomi, N., Kühl, M. & Kestler, H. A. Concepts in Boolean network modeling: What do they all mean? Comput. Struct. Biotechnol. J. 18, 571–582 (2020).

66. R: The R Project for Statistical Computing. Available at: https://www.r-project.org/. (Accessed: 28th February 2023)

67. seaborn: statistical data visualization — seaborn 0.11.1 documentation. Available at: https://seaborn.pydata.org/. x(Accessed: 29th June 2021)

